# Cellfm-datasets: A Unified Data Infrastructure for Single-Cell and Spatial Transcriptomics Foundation Model Pretraining

**DOI:** 10.64898/2026.06.11.731508

**Authors:** Liluojing Zhang, Jiangshuan Pang, Jing Yan, Wangyang Tang, Yiting Deng, Youzhe He

## Abstract

Large-scale cell foundation models are increasingly limited not only by model architecture, but also by the data infrastructure required to repeatedly sample sparse transcriptomic profiles from out-of-core cohorts. AnnData/H5AD has become a standard exchange format for single-cell and spatial omics analysis, yet its HDF5-backed layout is not designed for high-frequency random mini-batch loading under multi-worker and distributed pretraining. We present Cellfm-datasets, a data infrastructure artifact that converts H5AD cohorts into a self-describing compressed sparse row (CSR) memmap layout and exposes the resulting corpus through Hugging Face Dataset and IterableDataset interfaces. The artifact stores a shared gene vocabulary, per-sample metadata, optional spatial coordinates, observation metadata, manifests, and checksums, and reconstructs sparse cell or group records at runtime without dense expansion. A unified sampling abstraction supports random-cell groups, manifest-defined biological regions, and coordinate-based spatial blocks, with deterministic sharding across distributed ranks and data-loader workers. Spatial demonstrations on P14 mouse brain transcriptomics sections illustrate region- and block-level sampling over real anatomical structures. In controlled benchmarks on a public heterogeneous ModelScope scRNA-seq subset, Cellfm-datasets reached 60,571 ± 1,734 samples/s in single-core random loading, scaled to approximately 160,000 samples/s with eight workers, and maintained near-constant process-private memory while reading up to one million cells. By moving sparse single-cell and spatial corpora from model-specific loader code into reusable, validated, and framework-native dataset artifacts, this design may reduce the engineering burden of reproducible cell foundation model pretraining and make repeated training runs, model comparisons, and mixed-modality data reuse easier to standardize.

**Code availability:** https://github.com/PangJiangShuan/cellfm-datasets

## 1 Introduction

Single-cell RNA sequencing and spatial transcriptomics have moved computational biology from study-scale matrices toward atlas-scale corpora. Community resources such as the Human Cell Atlas [1], Tabula Sapiens [2], CZ CELLxGENE Discover [3], and Tahoe-100M [4] are large, heterogeneous, and increasingly reused across studies. Spatial assays, including Spatial Transcriptomics/Visium [5], Slide-seqV2 [6], and MERFISH [7], add coordinate systems and tissue context to expression profiles. Together, these resources have supported cell foundation models such as scBERT [8], Geneformer [9], scGPT [10], scFoundation [11], CellFM [12], and Nicheformer [13].

The training workload follows the broader transformer pretraining paradigm [14]: large corpora are sampled repeatedly, shuffled, and consumed by high-capacity sequence models. In biological settings, the data layer must also preserve sparse expression, gene-index consistency, cell annotations, and spatial coordinates while supporting multi-worker and distributed loading. Because the same cohort is often reused across model versions, hyperparameter sweeps, and distributed launches, data handling becomes an infrastructure problem rather than a minor input pipeline.

The current ecosystem is only partly organized around this infrastructure view. AnnData/H5AD [15], the de facto single-cell exchange format, is built on HDF5 [16, 17]. It remains highly interoperable for analysis, but high-frequency random row access under multi-worker pretraining is not its primary access pattern. As a result, many cell foundation model repositories include project-specific conversion scripts or custom PyTorch datasets. Such loaders may be adequate for a single study, but reuse, auditability, checksums, validation, and distributed sharding are usually tied to model-specific code. The limitation becomes more visible as pretraining data begins to mix dissociated single-cell collections with spatial transcriptomics.

Recent tools address parts of the problem. AnnLoader provides a PyTorch data-loader interface for AnnData objects [18]. scDataset combines block sampling with batched fetching over AnnData files for on-disk training [19]. BioNeMo-SCDL offers PyTorch-compatible single-cell loading over NumPy memory-mapped storage [20]. annbatch, a recent AnnData-native loader, supports out-of-core mini-batches for terabyte-scale biological training workloads [21]. These systems form relevant baselines and related infrastructure; among the compared baselines, to our knowledge, none provides one storage protocol for both coordinate-free single-cell data and coordinate-bearing spatial transcriptomics, or a native Hugging Face dataset interface for biological pretraining data [22].

We introduce Cellfm-datasets, a data infrastructure package for this training setting. Its design keeps expression sparse from conversion to batch assembly, stores single-cell and spatial transcriptomic samples under one protocol, and treats sampled cell groups as records rather than incidental loader outputs. Converted corpora are self-describing directories with manifests, checksums, and validation utilities, so assumptions about gene order, coordinates, and metadata are not hidden in model code. At runtime, the package exposes records through Hugging Face `Dataset/IterableDataset` objects and PyTorch data loaders [23].

### Contributions

This work makes five contributions: (i) a unified CSR-memmap storage protocol for H5AD cohorts that supports single-cell and spatial transcriptomics through the same directory layout, shared gene vocabulary, sample manifest, optional `obs.parquet`, optional coordinates, and checksum manifests; (ii) a runtime for memmap-backed sparse row access that reconstructs cells and cell groups from sparse row slices without dense expansion of high-dimensional expression vectors; (iii) a cell-group abstraction with random-cell, manifest-region, and spatial-block samplers for generic single-cell pretraining, annotated spatial regions, and coordinate-local tissue neighborhoods; (iv) Hugging Face `Dataset/IterableDataset` adapters with deterministic sharding across distributed ranks and data-loader workers, bridging cell omics data to the foundation-model data ecosystem; and (v) a software artifact with CLI conversion, validation, checksum, benchmark export, optional dependency groups, examples, and tests.

## 2 Related Work

### 2.1 Cell Foundation Models

Cell foundation models transfer the pretraining paradigm from language and vision to transcriptomic profiles. Representative systems include scBERT [8], Geneformer [9], scGPT [10], scFoundation [11], CellFM [12], and Nicheformer [13]. These models differ architecturally, but all depend on robust out-of-core sampling over sparse, heterogeneous, and repeatedly reused training corpora.

The data abstraction used by these models is also moving beyond one-cell-at-a-time retrieval. A training example may contain a single cell, a condition-specific set of cells, a local tissue neighborhood, or a region supplied by upstream biological annotation. As in document or image-patch pretraining, the modeling objective shapes the unit of sampling. A reusable data layer therefore needs a consistent way to assemble grouped sparse expression, coordinates, and metadata.

### 2.2 Data Formats for Single-Cell and Spatial Omics

AnnData and H5AD are central to the Python single-cell ecosystem because they package expression matrices, cell annotations, gene annotations, embeddings, and unstructured metadata in one interoperable object [15]. Scanpy [24], Bioconductor workflows [25], Seurat [26], scvitools [27], and SpatialData [28] all rely on stable containers for analysis and exchange. This work addresses a complementary need: turning H5AD cohorts into a training-oriented representation for row-random access during foundation model pretraining.

Sparse matrix formats are a natural fit for transcriptomics because most cell-by-gene entries are zero. CSR stores row pointers, column indices, and non-zero values, making retrieval of all expressed genes in one cell proportional to the number of non-zero entries rather than to the full vocabulary size. SciPy provides the standard CSR abstraction in the scientific Python stack [29], while NumPy supplies memory-mapped arrays that can be backed by the operating system page cache [30]. More general array storage systems, including TileDB [31], further motivate separating logical analysis objects from physical storage optimized for access patterns.

### 2.3 Spatial Transcriptomics Analysis

Spatial transcriptomic analysis introduces access patterns that are absent from conventional single-cell pretraining. Methods such as Tangram [32], cell2location [33], and SpaGCN [34] use tissue coordinates, cell-type mixtures, or histology to infer spatial organization. These settings motivate loaders that preserve local neighborhoods or anatomical regions instead of treating all cells as independent draws. Cellfm-datasets follows this view by representing each sampled group as a record over a shared sparse cell store.

### 2.4 Training Data Loaders

AnnLoader exposes AnnData collections through PyTorch data-loading utilities and converts selected fields to tensors [18]. scDataset uses block sampling and batched fetching to balance randomness and I/O locality over on-disk data [19]. BioNeMo-SCDL converts single-cell data into NumPy memory-mapped storage and provides PyTorch-compatible dataset classes [20]. annbatch adds AnnData-native out-of-core batching for large biological training pipelines [21]. These tools make clear that data loading can limit training throughput, although their interfaces remain primarily PyTorch-or AnnData-centered.

Hugging Face `datasets` provides a widely used abstraction for foundation-model data, including map-style datasets, iterable streaming datasets, deterministic sharding, caching, and local materialization [22]. Cellfm-datasets brings this interface to cell omics by exposing converted single-cell and spatial corpora as dataset records while preserving sparse expression and biological metadata.

## 3 Methods

### 3.1 System Overview and Design Requirements

Cellfm-datasets is designed for repeated biological foundation model pretraining rather than one-off analysis scripts. A typical user converts a cohort once, validates it, reuses it across training runs, and streams it into local or distributed training code. The main requirements are: (i) persistent conversion from H5AD to a layout that is efficient for row-level reads; (ii) preservation of sparsity and gene-index consistency across samples; (iii) support for coordinate-free single-cell samples and spatial samples with 2D or 3D coordinates; (iv) cell-and group-level access patterns; (v) reproducible distributed sharding; and (vi) compatibility with Hugging Face and PyTorch training code.

The package separates the data system into four layers. A storage layer defines the on-disk protocol; a runtime layer opens NumPy memmaps and reconstructs sparse cells or groups; a sampling layer produces modality-agnostic group specifications; and an adapter layer exports records as Hugging Face datasets and optional PyTorch collator inputs. With this separation, storage can be validated independently of model code, samplers can change without rewriting the storage format, and training code can consume standard dataset objects.

Fig. 1 summarizes the conversion-to-runtime pipeline and its training interfaces.

**Figure 1.**
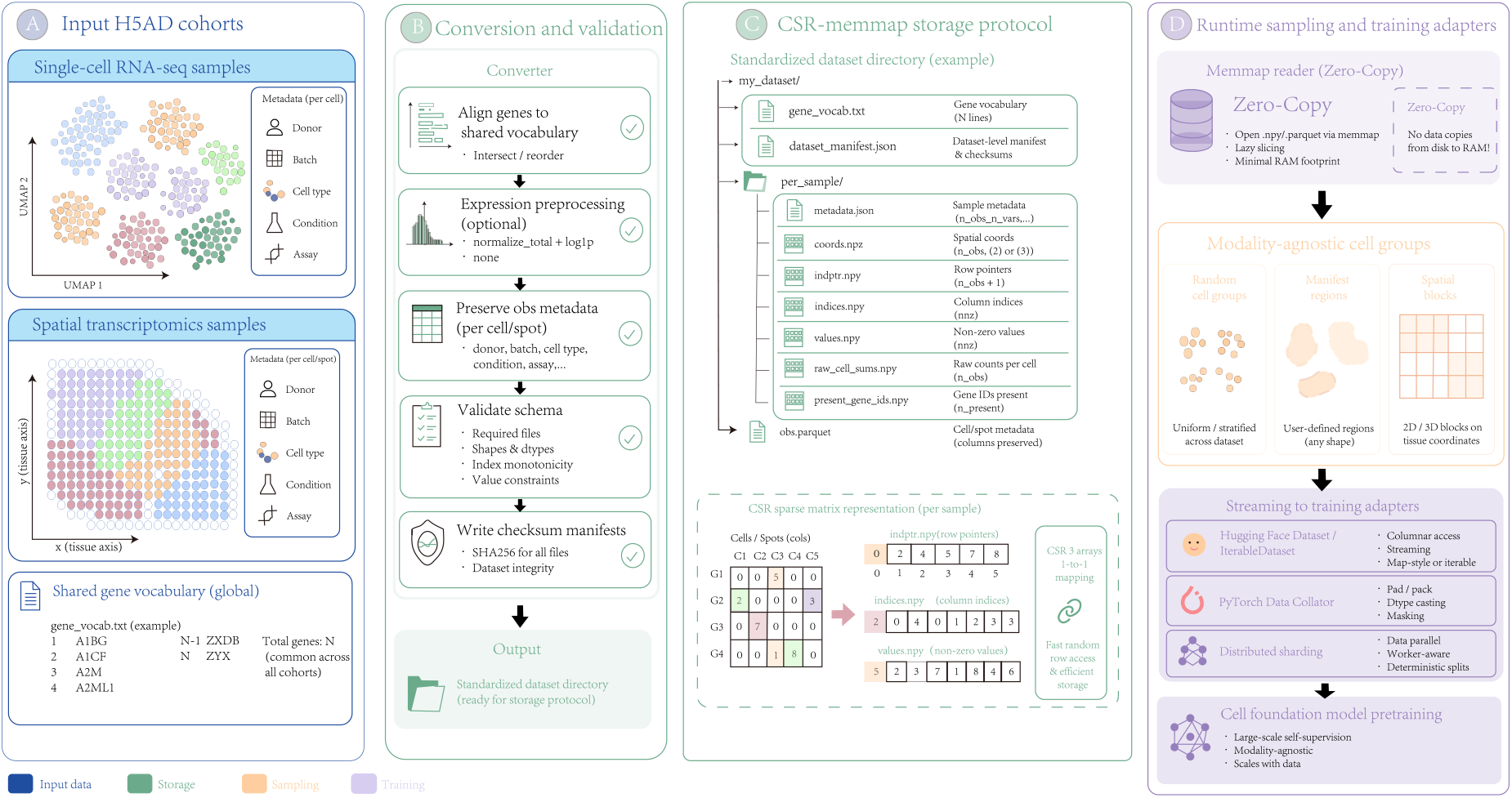
System architecture of Cellfm-datasets. H5AD cohorts from single-cell RNA-seq and spatial transcriptomics are converted by aligning genes to a shared vocabulary, applying optional expression preprocessing, preserving selected observation metadata, validating schema constraints, and writing checksums. The output is a standardized CSR-memmap directory. At runtime, readers perform memmap-backed sparse row access without dense expansion, samplers generate random cell groups, manifest-defined regions, or spatial blocks, and adapters expose the data to Hugging Face and PyTorch training workflows.

### 3.2 Canonical CSR-Memmap Layout

Each converted corpus is a directory with a root-level `gene_vocab.txt`, a `dataset_manifest. json`, and one subdirectory per sample. A sample directory stores three CSR arrays, `indptr.npy, indices.npy`, and `values.npy`; optional spatial coordinates in `coords.npy`; per-cell count summaries in `raw_cell_sums.npy`; present gene IDs; observation names; optional `obs.parquet`; and `metadata.json`. The integer gene IDs in `indices.npy` follow the shared vocabulary order, so all samples use the same gene coordinate system after conversion.

The same layout represents both single-cell and spatial transcriptomics. Spatial samples store a `coords.npy` matrix with two or three coordinate axes. Coordinate-free single-cell samples store a valid array of shape (*n*_cells_, 0) and set `coord_dim` to zero. Thus, every sample exposes expression, coordinates, metadata, and row-level accessors, even when the coordinate component is empty. During conversion, genes are aligned to a shared vocabulary. Expression can either be stored after total-count normalization followed by `log1p`, or preserved without additional transformation. Sample metadata records the expression transform, number of cells, number of genes, non-zero count, coordinate dimension, coordinate axis names, retained and dropped genes, and stored observation columns. Validation checks required files, array shapes, CSR invariants, manifest consistency, and optional region-manifest examples.

### 3.3 Random-Access Runtime

The runtime uses a sample reader and a cohort reader. For a cell index *i*, the row interval is obtained from `indptr[i]` and `indptr[i+1]` . The runtime then slices `indices` and `values` only over that interval. Thus, the I/O cost for one cell is *O*(nnz_*i*_) rather than *O*(*G*), where *G* is the number of genes. For group construction, the runtime fetches the selected row intervals, builds a local pointer array `cell_ptr`, and concatenates the corresponding gene indices and values. It does not densify the expression matrix.

All core arrays are opened with NumPy `mmap mode=“r”`. Initialization therefore avoids loading the full cohort into the process heap, and multiple worker processes can read the same file-backed pages through the operating system page cache. Observation metadata are stored separately in `obs.parquet` and loaded lazily only when a caller requests metadata fields.

### 3.4 Cell Group Abstraction and Sampling

The package treats a cell group as the primary training unit. A group is defined by a sample identifier, a group identifier, a group type, and a set of source cell indices; expression, coordinates, and metadata are reconstructed at runtime. The returned record contains local sparse expression arrays, raw and optionally normalized coordinates, source cell indices, optional observation metadata, anchor coordinates, block bounds, and sampling metadata. Through this abstraction, single-cell random sampling and spatial neighborhood sampling share the same downstream interface. It also matches pretraining settings in which a model consumes a matrix or local context rather than a single isolated cell.

Three samplers are currently supported:

*RandomCellSampler* samples fixed-size groups from one or more samples, using uniform or cell-count-proportional sample weighting and optional observation-column stratification. It is suited for generic single-cell pretraining.

*ManifestRegionSampler* reconstructs groups from an external region manifest with sample IDs, region IDs, and source cell indices. The region manifest can come from annotation, segmentation, anatomical labeling, or atlas pipelines. Keeping the manifest outside the core storage format allows different region definitions to be evaluated over the same converted dataset. If the manifest includes anchor-coordinate fields or region metadata, the runtime forwards them into the group record.

We demonstrate this sampler on a postnatal day 14 (P14) mouse brain development spatial transcriptomics section. This stage captures neurodevelopmental tissue organization and contains fine anatomical partitions such as cortex and hippocampus. The data include anatomical annotation boundaries (`scc_anno`) and molecular clustering labels (`louvain`). Fig. 2(a)–(d) shows the P14 tissue coordinates, anatomical-region manifest, region sizes, and anchor coordinates; Fig. 2(e)–(j) shows six manifest-defined batches. Unlike purely random cell sampling or rectangular grid cropping, *ManifestRegionSampler* can use these biological priors to assemble an irregular anatomical structure, such as a complete hippocampal-shaped region, into one training group while preserving the original coordinate topology.

**Figure 2.**
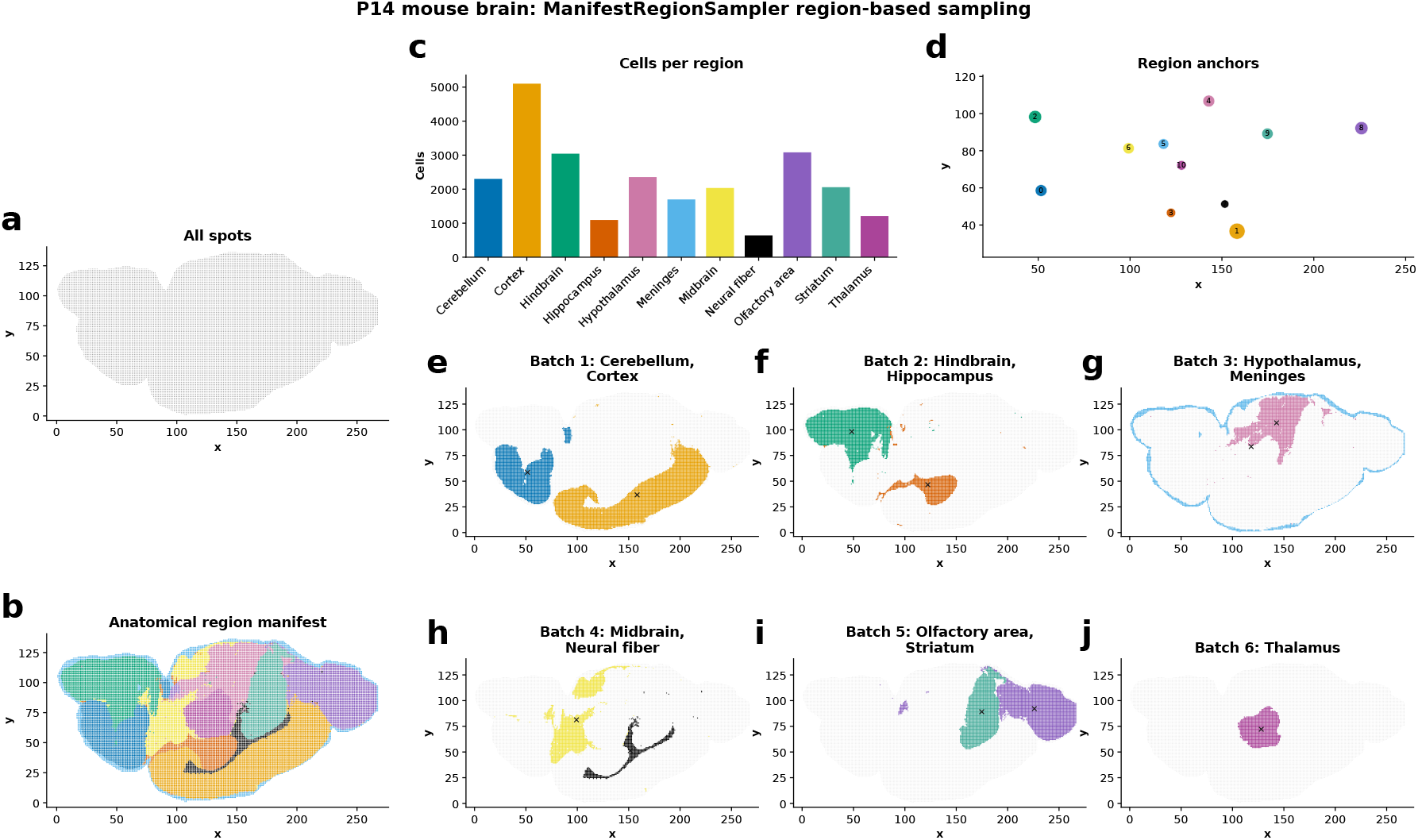
Manifest-region sampling on P14 mouse brain development spatial transcriptomics. (a) All spatial spots. (b) `scc_anno`-derived anatomical region manifest. (c) Cells per anatomical region. (d) Region anchors. (e)–(j) Six manifest-defined batches: cerebellum/cortex, hindbrain/hippocampus, hypothalamus/meninges, midbrain/neural fiber, olfactory area/striatum, and thalamus. The sampler constructs irregular biological regions while preserving source coordinates and sparse expression.

*SpatialBlockSampler* generates groups from coordinate-space blocks or from a custom block manifest. In grid mode, it computes coordinate ranges for each spatial sample, tiles the coordinate space using a user-defined block shape and stride, filters blocks with too few cells, and optionally downsamples dense blocks to a maximum cell count. In manifest mode, blocks may be supplied by external tissue segmentation or image-analysis pipelines. Fig. 3(a)–(e) compares all spots, regular grid blocks, custom block manifests, and their corresponding block-size distributions; Fig. 3(f)–(k) shows coordinate-local batches generated from selected blocks. This mode is intended for spatial pretraining where local tissue neighborhoods should remain visible to the model.

**Figure 3.**
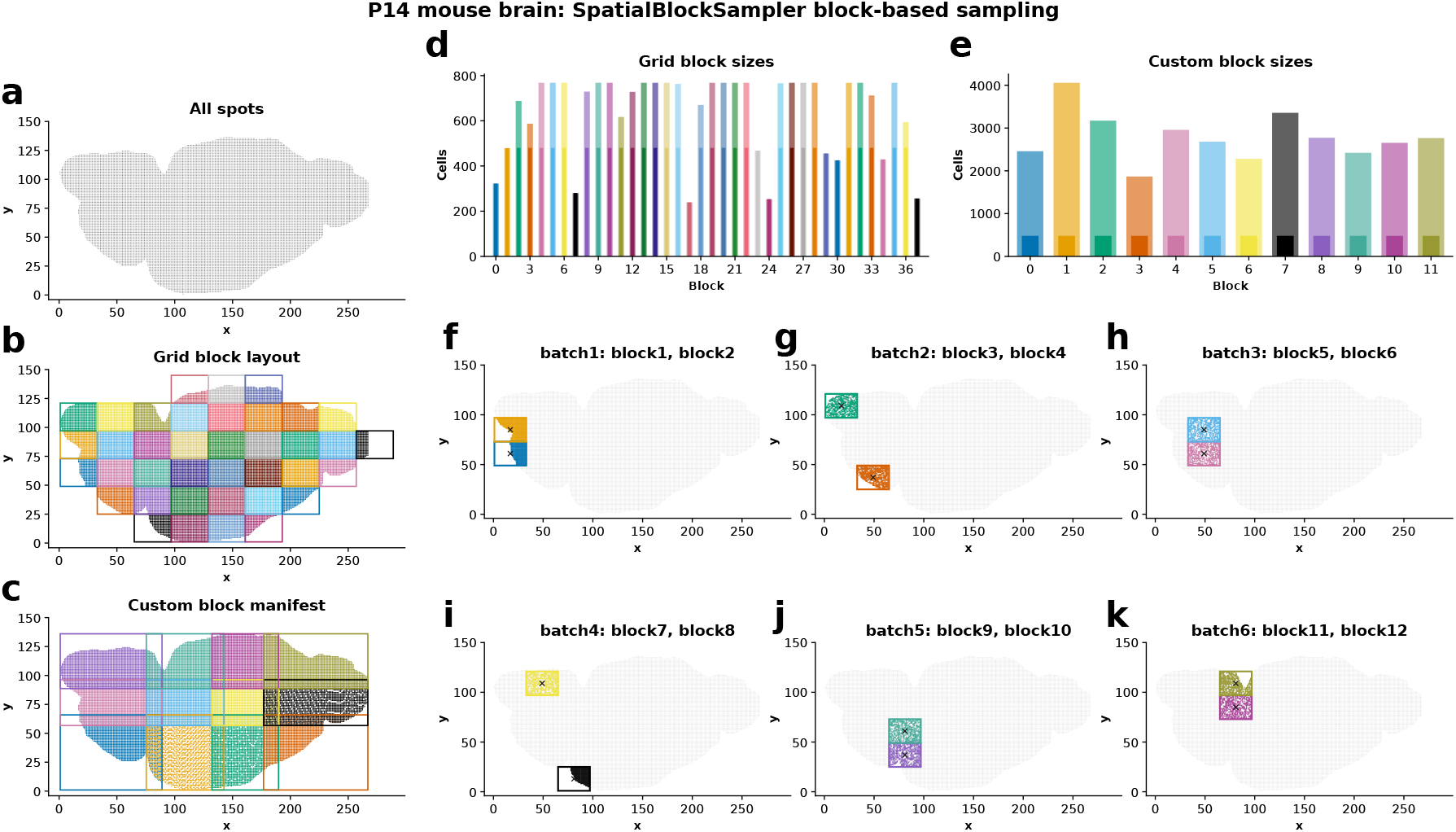
Spatial-block sampling on P14 mouse brain development spatial transcriptomics. (a) All spatial spots. (b) Regular grid block layout. (c) Custom block manifest aligned to tissue geometry. (d) Cell counts for grid blocks. (e) Cell counts for custom blocks. (f)–(k) Six coordinate-local batches from block pairs 1–2, 3–4, 5–6, 7–8, 9–10, and 11–12. The sampler generates local tissue-neighborhood groups for spatial pretraining.

### 3.5 Hugging Face and Distributed Integration

Cellfm-datasets exposes cell-, group-, and region-level Hugging Face adapters. Each adapter returns either a map-style `Dataset` or streaming `IterableDataset`. Cell-level records contain sample ID, cell index, observation name, coordinates, raw cell sum, gene indices, and gene values. Group-level records additionally contain group ID, group type, source cell indices, a local CSR pointer array, optional anchor coordinates, block bounds, and sampling metadata. The public API includes `load_hf_cell_dataset`, `load_hf_group_dataset`, and `load_hf_region_dataset`, allowing users to select the granularity that matches their model.

Distributed execution is handled by a runtime context that records rank, world size, worker ID, number of workers, and epoch. The runtime performs stride-based sharding over examples or sampler group indices, making the stream non-overlapping across ranks and data-loader workers. Random samplers use epoch-aware seed offsets so that multi-epoch training remains reproducible while changing the sampled groups.

### 3.6 Engineering and Release Tooling

Cellfm-datasets is provided as a software artifact with a small required dependency set and optional extras for conversion, Hugging Face integration, benchmarking, plotting, Torch collation, and development. The command-line interface supports H5AD conversion, dataset inspection, schema validation, checksum manifest creation and verification, benchmark data generation, and local export of selected splits. These utilities make the dataset artifact reusable, testable, and distributable independently of any specific model repository.

## 4 Experiments

### 4.1 Baselines

We benchmarked Cellfm-datasets against AnnLoader, scDataset, and BioNeMo-SCDL because they represent practical alternatives for training-oriented access to AnnData-derived single-cell data. *AnnLoader* stays closest to the native AnnData ecosystem by exposing AnnData objects through PyTorch-style loading utilities. *scDataset* targets large-scale single-cell training with block sampling and batched fetching over on-disk data. *BioNeMo-SCDL* represents PyTorch-compatible memory-mapped single-cell storage. Because Cellfm-datasets is positioned as a reusable data infrastructure layer, the evaluation considers sparse storage, worker scaling, memory behavior, sampling diversity, spatial support, and Hugging Face integration rather than raw speed alone.

### 4.2 Benchmark Setup

The spatial demonstrations in Fig. 2 and Fig. 3 used P14 mouse brain development spatial transcriptomics data from the single-cell spatial transcriptomic atlas of the whole mouse brain [35]; the associated data accession is DOI: 10.12412/BSDC.1699433096.20001. This dataset provides anatomical annotations and molecular clusters for region- and block-level access over real tissue structures. The single-cell benchmarks used the public ModelScope scRNA-seq collection [36]: 40 human H5AD files with approximately 1.13 million cells across tonsil, breast, colon, and bone marrow, including healthy samples and clinical disease samples such as Alzheimer’s disease, breast cancer, and COVID-19. Its batch variation, gene-dimensional differences, and cell-type labels make it a useful testbed for heterogeneous data loading and random-throughput behavior.

All tools were evaluated in the same CPU-only workstation setting with local SSD/NVMe storage, identical input cohorts, and matched batch-size, warmup, and measurement schedules. Throughput and memory benchmarks were repeated five times; plots report the mean with standard-deviation error bars where shown. We report samples per second, peak private dirty memory, and batch-level Shannon entropy.

We evaluated four settings: single-core random loading throughput, multi-worker throughput, memory growth while reading increasing numbers of cells, and batch-level sampling randomness.

## 5 Results

### 5.1 Single-Core Loading Throughput

Fig. 4 reports single-core random data loading throughput with batch size 64 and no worker processes. This experiment isolated the cost of reading cells from the underlying storage format. Across five runs, Cellfm-datasets achieved a mean throughput of 60,571 samples/s, compared with 16,489 samples/s for BioNeMo-SCDL, 1,615 samples/s for scDataset, and 266 samples/s for AnnLoader.

**Figure 4.**
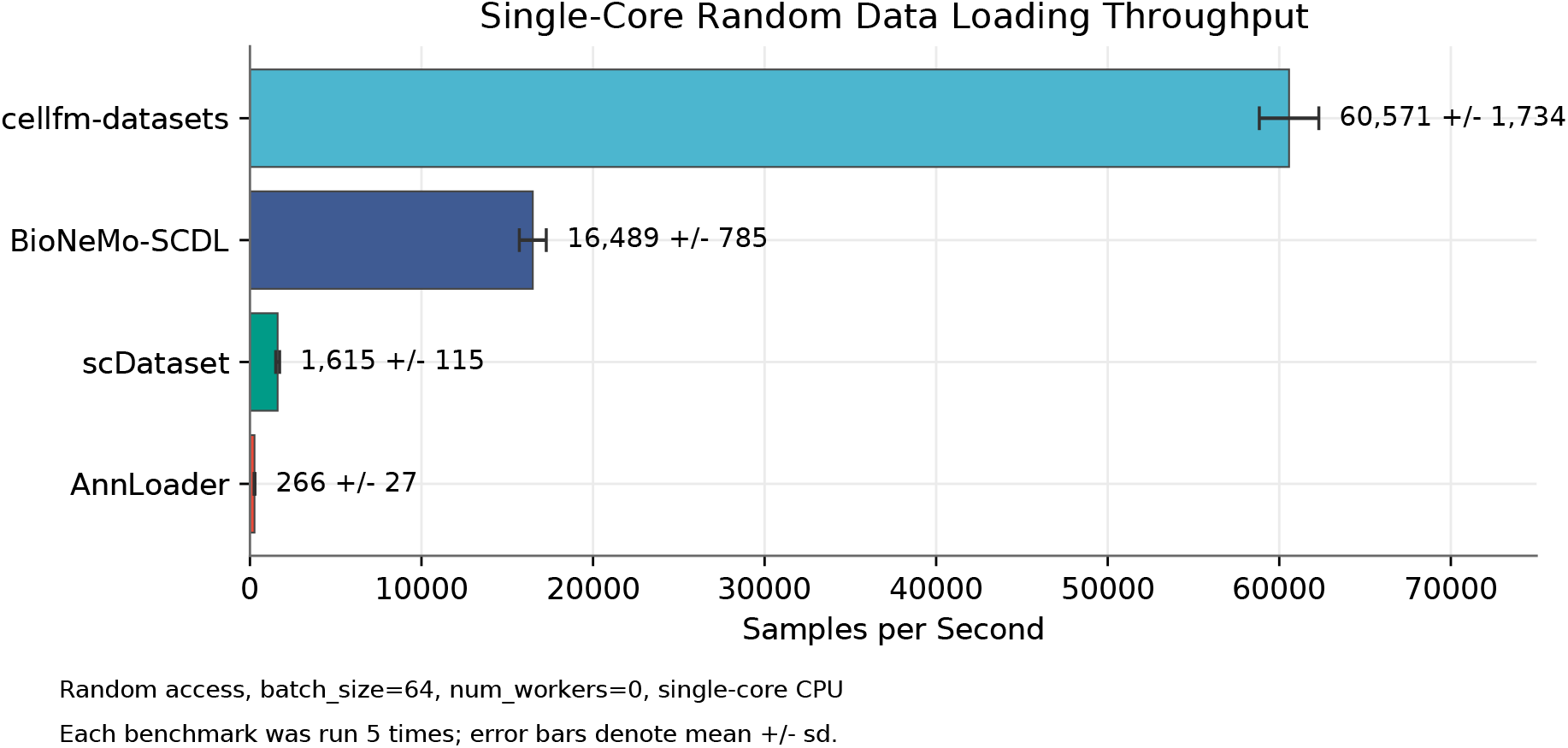
Single-core random loading throughput on the heterogeneous ModelScope scRNA-seq subset. The CPU-only benchmark uses local SSD/NVMe storage, random access, batch size 64, and no worker processes. Bars report the mean of five runs, and error bars denote standard deviation.

The gap is consistent with reading only non-zero CSR row slices under a random-access workload. AnnLoader incurred HDF5-backed access through the AnnData stack. scDataset improved direct H5AD access through block-aware fetching, although this particular test stressed true random reads. BioNeMo-SCDL benefited from memmap storage, but dense or less sparse-native reads still moved more data per cell than CSR row slicing. These measurements should be read as workload-specific rather than as a universal ranking across all loading regimes.

### 5.2 Worker Scaling

Fig. 5 evaluates throughput as the number of data-loader workers increased. The setting approximated the input side of large-model training, where each GPU process may launch multiple workers to prepare mini-batches. AnnLoader failed or was unsupported for the tested worker settings and is marked accordingly in the figure. The remaining three tools were compared over one, two, four, and eight workers.

**Figure 5.**
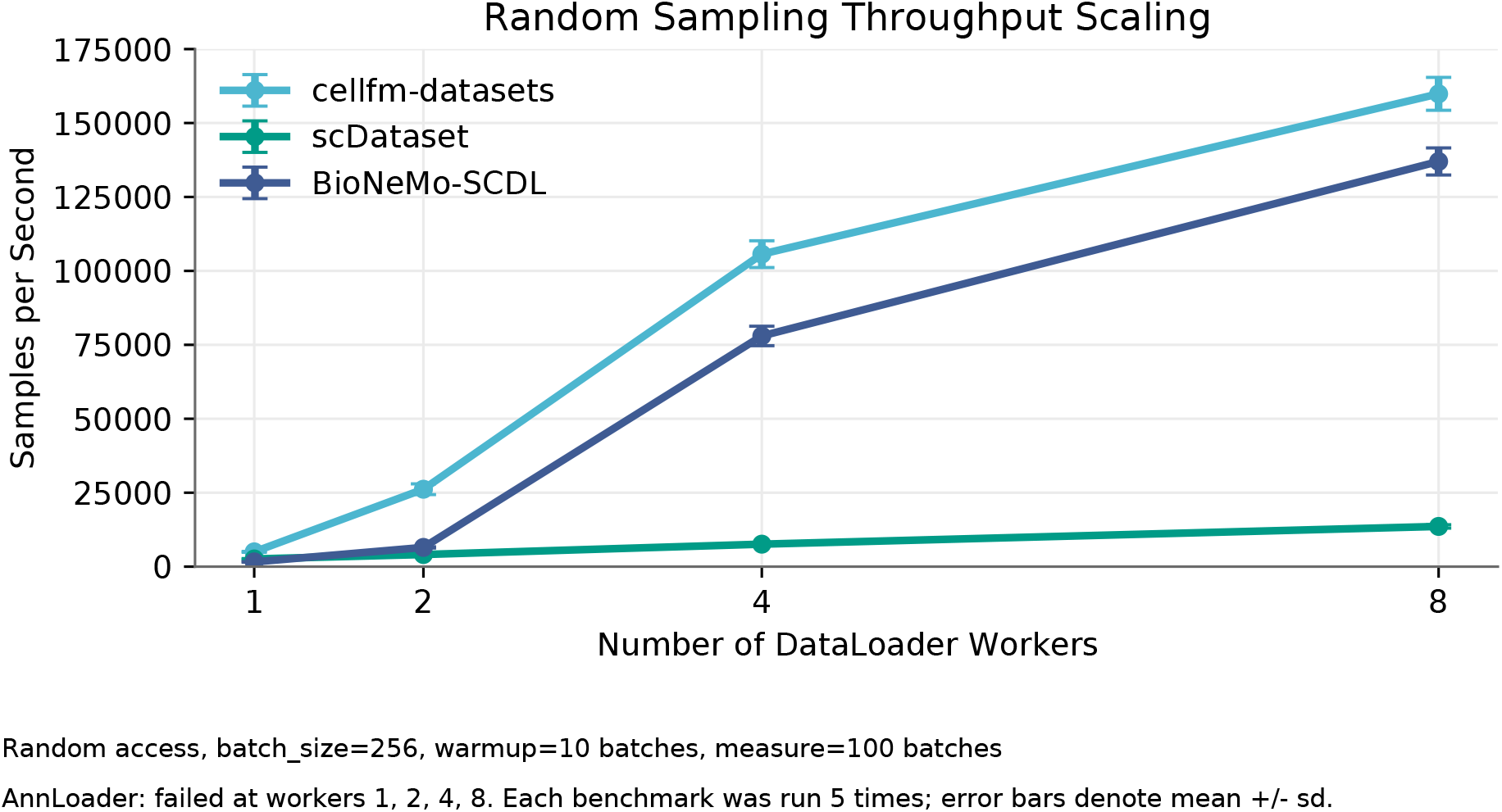
Random sampling throughput scaling with data-loader workers. The benchmark uses random access, batch size 256, 10 warmup batches, and 100 measured batches. Markers report the mean of five runs, and error bars denote standard deviation.

Cellfm-datasets scaled from approximately 5,000 samples/s with one worker to approximately 160,000 samples/s with eight workers. BioNeMo-SCDL also benefited from memmap-based access but remained below Cellfm-datasets in this benchmark. scDataset improved with additional workers, though it remained limited under the random-access configuration. The observed scaling is consistent with sparse row-native reads reducing per-sample data movement and read-only memmap pages being shared by worker processes through the operating system page cache.

### 5.3 Memory Efficiency

Fig. 6 compares peak private dirty memory as the number of randomly read cells increased. We used private dirty memory rather than total resident memory because it better reflects memory uniquely owned by the process; shared file-backed pages from memmap readers should not be counted as private heap growth. AnnLoader grew approximately linearly and reached nearly 50 GiB at one million read cells in this benchmark. scDataset and BioNeMo-SCDL stayed near 4 GiB, while Cellfm-datasets remained close to the baseline with only small increases.

**Figure 6.**
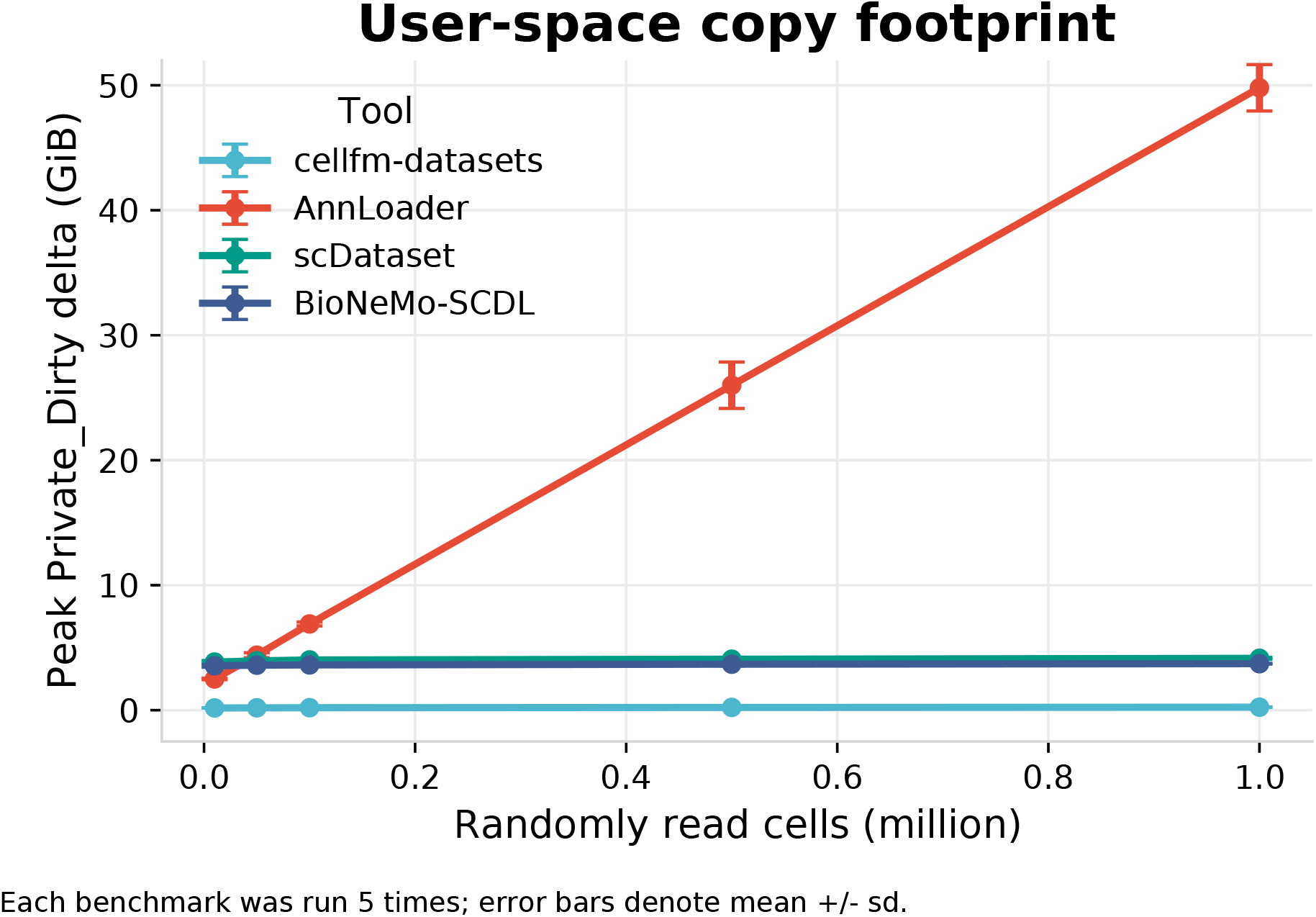
Peak process-private memory while randomly reading increasing numbers of cells. Markers report the mean of five runs, and error bars denote standard deviation. The private-dirty metric distinguishes dense in-memory loading from shared file-backed memmap access.

This behavior follows the CSR-memmap design. The private heap stores the current sparse slices, local pointer arrays, and small metadata objects; large expression arrays remain file-backed and are paged by the operating system. For foundation model pretraining, this memory profile is useful because CPU memory must also support preprocessing, data-loader workers, model input buffers, and distributed runtime processes.

### 5.4 Batch Diversity

Randomness and biological diversity affect self-supervised pretraining. A loader that repeatedly reads local blocks may reduce stochasticity even when it is fast. Fig. 7 reports the distribution of batch-level cell-type Shannon entropy as batch size increased; higher entropy indicates more diverse cell-type composition within a batch.

**Figure 7.**
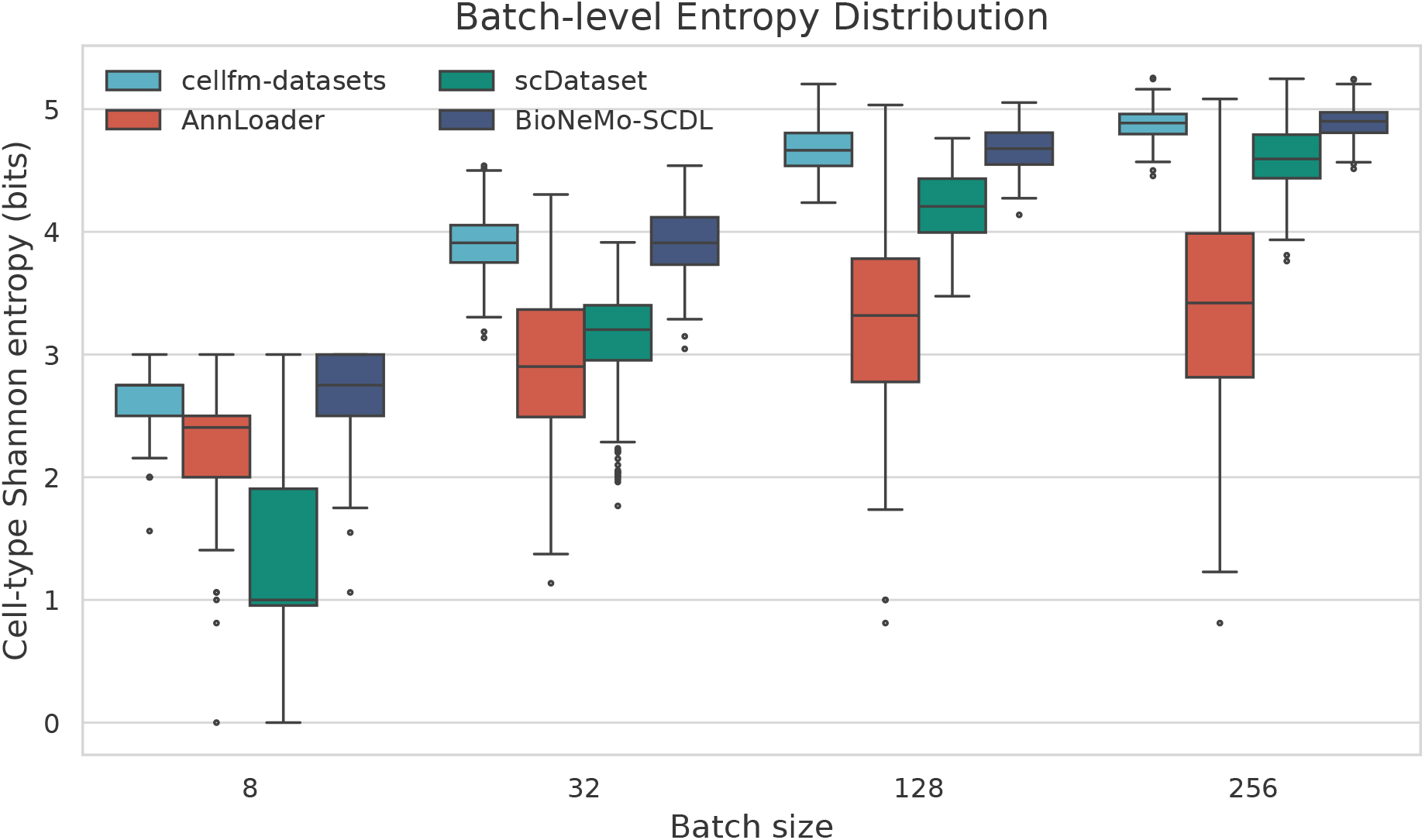
Batch-level cell-type Shannon entropy across batch sizes. Higher entropy indicates more diverse cell-type composition within a batch. This plot is intended to complement throughput benchmarks by measuring the sampling behavior seen by the model.

Cellfm-datasets and BioNeMo-SCDL maintained high entropy at larger batch sizes under true random access. scDataset improved with batch size but reflected the locality introduced by block sampling. AnnLoader showed larger variance. These results suggest that Cellfm-datasets can support high throughput without fixing one sampling regime: users may choose fully random groups for global mixing, manifest-region groups for biologically annotated spatial regions, or spatial blocks for local tissue context according to the pretraining objective.

## 6 Discussion

Cellfm-datasets occupies a different point in the design space from existing loaders: it combines persistent sparse storage, manifest-backed and coordinate-local group sampling, and standard foundation-model data interfaces. The P14 mouse brain examples show why a unified protocol is useful. Anatomical regions are often irregular, yet they can define meaningful training groups. Encoding coordinate-free cells and coordinate-bearing spatial samples in the same layout moves these choices into dataset metadata rather than ad hoc model branches.

The reusable artifact is also intended to make assumptions auditable. Gene order, metadata fields, checksums, validation state, and sharding behavior are recorded outside the model repository. Its Hugging Face interface further avoids repeated implementation of streaming, caching, and worker partitioning while preserving sparse expression.

Table 1 and Table 2 summarize representative runtime settings and qualitative differences among the compared tools.

**Table 1.** Representative Cellfm-datasets throughput and sampling diversity.

| Batch | Workers | Samples/s | Entropy |
| --- | --- | --- | --- |
| 4 | 1 | 3,840 | 2.10 |
|  | 2 | 8,920 | 2.12 |
|  | 4 | 20,740 | 2.16 |
|  | 8 | 36,580 | 2.18 |
| 16 | 1 | 4,320 | 3.18 |
|  | 2 | 14,860 | 3.20 |
|  | 4 | 44,530 | 3.22 |
|  | 8 | 75,240 | 3.24 |
| 64 | 1 | 4,760 | 4.33 |
|  | 2 | 22,140 | 4.35 |
|  | 4 | 82,600 | 4.37 |
|  | 8 | 126,400 | 4.39 |
| 256 | 1 | 5,002 | 4.92 |
|  | 2 | 26,175 | 4.94 |
|  | 4 | 105,656 | 4.97 |
|  | 8 | 159,870 | 4.90 |

**Table 2.** Functional comparison of data-loading tools. HF denotes native Hugging Face interfaces.

| Tool | Storage | Modality | Group / sampling | HF | Shard |
| --- | --- | --- | --- | --- | --- |
| AnnLoader | H5AD / AnnData | SC | cell mini-batches | – | PyTorch |
| scDataset | H5AD block access | SC | block-aware fetching | – | PyTorch |
| BioNeMo-SCDL | NumPy memmap | SC | cell-level dataset | – | PyTorch |
| **Cellfm-datasets** | **CSR-memmap** | **SC+ST** | **random, region, block groups** | **✓** | **rank + worker** |

The main trade-off is the offline conversion step. This cost is easiest to justify when a cohort is reused across training runs, model sizes, or distributed configurations; for small exploratory studies, direct AnnData access may remain simpler. The package therefore complements AnnData with a training-oriented representation rather than replacing it as an analysis object.

The current implementation targets transcriptomic matrices and optional coordinates; ATAC-seq, CITE-seq, imaging features, histology images, and complex multi-omic tensors remain future work. Variable-size spatial blocks require a custom collator or fixed-size sampling. Broader evaluations across file systems, storage devices, multi-node clusters, and sampler ablations are still needed.

## 7 Conclusion

We presented Cellfm-datasets, a sparse memmap data infrastructure artifact for biological foundation model pretraining. It converts H5AD cohorts into a canonical CSR-memmap layout, preserves single-cell and spatial transcriptomics under one protocol, reconstructs random or spatially structured cell groups, and exposes the data through Hugging Face and PyTorch-compatible interfaces.

Benchmarks against AnnLoader, scDataset, and BioNeMo-SCDL suggest that sparse row-native storage can provide high random-access throughput, worker scaling, low private memory usage, and competitive sampling diversity. Future work will extend the protocol to additional omics and imaging modalities, broaden atlas-scale evaluations, and develop richer collators for variable-size spatial and multi-modal cell groups.

